# Neutralizing antibody responses and cellular responses against severe acute respiratory syndrome coronavirus 2 omicron subvariant BA.5 after an mRNA severe acute respiratory syndrome coronavirus 2 vaccine dose in kidney transplant recipients

**DOI:** 10.1101/2023.08.02.551424

**Authors:** Keita Kawashiro, Rigel Suzuki, Takuto Nogimori, Naoya Iwahara, Takayuki Hirose, Kazufumi Okada, Takuya Yamamoto, Takasuke Fukuhara, Kiyohiko Hotta, Nobuo Shinohara

**Author notes:** These authors contributed equally to this work (Keita Kawashiro, Rigel Suzuki, Takuto Nogimori). These authors are corresponding authors. Correspondence authors: Takuya Yamamoto Ph.D., Laboratory of Precision Immunology, Center for Intractable Diseases and ImmunoGenomics, National Institutes of Biomedical Innovation, Health and Nutrition, 7-6-8, Saito-Asagi, Ibaraki City, Osaka, 567-0085, Japan; Takasuke Fukuhara M.D., Ph.D., Department of Microbiology and Immunology, Faculty of Medicine, Hokkaido University, Kita-ku, Kita-15 Nishi-7, Sapporo, Hokkaido, 060-8638, Japan. Kiyohiko Hotta M.D., Ph.D., Department of Urology, Hokkaido University Hospital, Kita-ku, Kita-14 Nishi-5, Sapporo, Hokkaido, 060-0814, Japan.

## Abstract

We examined the anti-severe acute respiratory syndrome coronavirus 2 (SARS-CoV-2) spike protein IgG antibody and neutralizing antibody titers and cellular immunity in 73 uninfected recipients and 17 uninfected healthy controls who received three doses of a coronavirus 2019 mRNA vaccine. Neutralizing antibody titers were evaluated using GFP-carrying recombinant SARS-CoV-2 with spike protein of B.1.1, omicron BA.1, or BA.5. For cellular immunity, peripheral blood mononuclear cells were stimulated with peptides corresponding to spike protein antigens of B.1.1, BA.1, and BA.5; spike-specific CD4/CD8 memory T cells were evaluated using intracellular cytokine staining. The median IgG antibody titers were 7.8 AU/mL in recipients and 143.0 AU/mL in healthy controls (p < 0.0001). Neutralizing antibody titers against all three viral variants were significantly lower in recipients (p < 0.0001). The number of spike-specific CD8 + memory T cells significantly decreased in recipients (p < 0.0001). Twenty recipients and seven healthy controls additionally received a bivalent omicron-containing booster vaccine, and IgG antibody and neutralizing antibody titers increased in both groups; however, the increase was significantly lower in recipients. Recipients did not gain sufficient immunity with a third dose of vaccine, suggesting a need to explore methods other than vaccines.

## 1. Introduction

Severe acute respiratory syndrome coronavirus 2 (SARS-CoV-2) infection started in Wuhan, China, in 2019 and has since become a pandemic. As of July 24, 2023, SARS-CoV-2 has infected more than 768 million people worldwide, causing more than 6.9 million deaths. The mortality rate of SARS-CoV-2 before vaccination has been reported to be higher in kidney transplant (KTX) recipients than in healthy controls (20–28% vs. 1–5%).^1^ Even after vaccination, KTX recipients have been reported to have a lower antibody acquisition rate, a higher infection, rate, and severe disease.^2, 3^ The antibody acquisition rate in KTX recipients after the third dose of the vaccine has been shown to be 53% and in healthy controls to be almost 100%, and KTX recipients have more severe SARS-CoV-2 infection than healthy controls.^3–5^ Furthermore, KTX recipients have been shown to have lower neutralizing antibody titers and cellular responses against B.1.1 (WT), B.1.617.2 (Delta), and B.1.1.529 (Omicron BA.1) after SARS-CoV-2 vaccination than healthy controls.^6–8^ However, to our knowledge, few studies have evaluated neutralizing antibody responses and cellular responses against Omicron BA.5 in KTX recipients. In this study, we investigated anti-SARS-CoV-2 spike protein IgG and neutralizing antibody titers and cellular responses against the Omicron subvariant BA.5 in KTX recipients after the third dose of an mRNA vaccine. Furthermore, we evaluated the effects of bivalent Omicron-containing booster vaccines on KTX recipients.

## 2. Materials and Methods

### 2.1 Study design and patients

We retrospectively reviewed the data of 403 recipients who underwent kidney transplantation at Hokkaido University between 1996 and 2021. Our inclusion criteria consisted of patients who were administered up to third mRNA vaccines after kidney transplantation. Conversely, the following criteria were used to exclude patients from the study: 1) loss of graft function and 2) SARS-CoV-2 infection prior to the study. Thus, 73 patients were included in the final analysis. As healthy controls, we reviewed the data from seven kidney donors who underwent nephrectomy at Hokkaido University Hospital and 10 clinical staffs at Hokkaido University. Serum was collected from the recipients and controls after the third doses of the SARS-CoV-2 vaccine. The study protocol was approved by the Institutional Review Board of Hokkaido University Hospital (approval number: CRB1180001). This study was conducted in accordance with the principles of the Declaration of Helsinki, 1996. Written informed consent was obtained from all the patients and healthy volunteers.

### 2.2 Measurement of anti-SARS-CoV-2 spike protein IgG antibody

Serum samples were collected, and anti-SARS-CoV-2 spike protein IgG antibodies were measured using a fully automated chemiluminescent enzyme immunoassay (CLEIA). The CLEIA was performed using SARS-CoV-2 S-IgG (IB) reagents (Fujirebio, Tokyo, Japan) and a Lumipulse L2400 system (Fujirebio). Anti-SARS-CoV-2 spike protein IgG antibody titers were measured at SRL (Tokyo, Japan). The cutoff value was defined as 1.0 AU/mL.

### 2.3 Cell culture

TMPRSS2-expressing Vero E6 (VeroE6/TMPRSS2) cells were obtained from the Japanese Collection of Research Bioresources Cell Bank (JCRB1819) and cultured in low-glucose Dulbecco’s modified Eagle’s medium (DMEM; Sigma-Aldrich, St. Louis, MO, USA) containing 10% fetal bovine serum (FBS) (Biowest, Bradenton, France) and G418 (Nacalai Tesque, Kyoto, Japan). HEK293-C34 cells were gifted by Y Matsuura at Osaka University and maintained in high-glucose DMEM (Nacalai Tesque) containing 10% FBS and 10 μg/mL blasticidin (solution) (InvivoGen, California, USA), and the exogenous expression of ACE2 and TMPRSS2 was induced by the addition of doxycycline hydrochloride (1 μg/mL) (Sigma-Aldrich). All the above cells were cultured at 37°C under 5% CO_2_.

### 2.4 SARS-CoV-2 reverse genetics

Recombinant SARS-CoV-2 was generated using a circular polymerase extension reaction (CPER) as previously described.^9^ Briefly, nine DNA fragments encoding the partial genome of SARS-CoV-2 (strain 2019-nCoV/Japan/TY/WK-521/2020, GISAID ID: EPI_ISL_408667) were amplified with polymerase chain reaction (PCR) using PrimeSTAR GXL DNA polymerase (Takara Bio Inc., Shiga, Japan). The corresponding SARS-CoV-2 genomic regions, PCR templates, and primers used for this procedure are summarized in Table S1. A linker fragment encompassing the hepatitis delta virus ribozyme, bovine growth hormone poly A signal, and cytomegalovirus promoter was also prepared using PCR. Ten obtained DNA fragments were mixed and used for CPER.^9^ To prepare green fluorescent protein (GFP)-expressing replication-competent recombinant SARS-CoV-2, we used fragment 9, in which the GFP gene was inserted into the ORF7a frame instead of the authentic F9 fragment.^9^ The rBA.1 S-GFP virus was gifted by K. Sato at Tokyo University.^10^ To prepare rBA.5 S-GFP virus, the fragment of viral genome that corresponds to the region of fragment 8 was subcloned from a BA.5 isolate (strain hCoV-19/Japan/TKYS14631/2022; GISAID ID: EPI_ISL_12812500). Nucleotide sequences were determined using a DNA sequencing service (Fasmac, Kanagawa, Japan), and sequence data were analyzed using ApE. To produce recombinant SARS-CoV-2 (seed virus), CPER products were transfected into HEK293-C34 cells using TransIT-LT1 (Takara) according to the manufacturer’s protocol. One day post-transfection, the culture medium was replaced with high-glucose DMEM (Nacalai Tesque) containing 2% FBS, 1% Penicillin-Streptomycin (PS), and 1μg/mL doxycycline. At 6–10 days post-transfection, the culture medium was harvested and centrifuged, and the supernatants were collected as seed viruses.

### 2.5 SARS-CoV-2 preparation and titration

The chimeric recombinant SARS-CoV-2 (rB.1.1 S-GFP, rBA.1 S-GFP and rBA.5 S-GFP) was amplified in Vero E6/TMPRSS2 cells, and the culture supernatants were harvested and stored at −80 until use. Infectious titers in the culture supernatants were determined using 50% tissue culture infective doses (TCID50). The TCID50 was calculated using the Reed-Muench method. The culture supernatants of the cells were inoculated onto VeroE6/TMPRSS2 cells in 96-well plates after serial 10-fold dilution with low-glucose DMEM containing 2% FBS and 1 mg/mL G418, and the infectious titers were determined 96 h post-infection. All experiments involving SARS-CoV-2 were performed in biosafety level-3 laboratories following standard biosafety protocols approved by Hokkaido University.

### 2.6 Neutralizing antibody titer assay

In each well, 7.5 × 10^3^ VeroE6/TMPRSS2 cells were seeded in 96-well plates and maintained in DMEM (high glucose) containing 10% FBS and 1% PS. The cells were then incubated overnight. On the following day, each serum sample was serially diluted three-fold in the culture medium, with a first dilution of 1:10 (final dilution range of 1:21,870). The diluted serum was incubated with 700 TCID50 of the chimeric recombinant SARS-CoV-2 at 37 in 5% CO_2_ for 1 h. Then, the mixture of chimeric recombinant SARS-CoV-2 and serum was added to VeroE6/TMPRSS2 cells in a 96-well plate. After 1 h, the cells were washed with DMEM (high glucose) containing 10% FBS and 1% PS. In the case of using GFP fluorescence, after incubating the plates at 37°C for 34–36 h, GFP fluorescence was detected using ECLIPSE Ts2 (Nikon, Tokyo, Japan). Then, the luminance of GFP was calculated using Image J. A GFP signal with a luminance value > 150 in one field of view was regarded as positive. The titer of the neutralizing antibody was defined as the minimum serum dilution at which the GFP signal was positive. The neutralizing antibody titer of each serum sample was defined as the average titer of the neutralizing antibody in triplicate assays. Data were plotted using GraphPad Prism 9 software (GraphPad Software, MA, USA).

### 2.7 Analysis of SARS-CoV-2 spike-specific T cells

Heparin-treated whole blood was drawn into Gebrauchsanleitung Leucosep (Greiner Bio-One) and processed to isolate peripheral blood mononuclear cells (PBMCs) according to the manufacturer’s instructions. Isolated PBMCs were cryopreserved in dimethyl sulfoxide (Sigma-Aldrich) and stored in −196°C liquid nitrogen until they were used in the assays. For analyzing SARS-CoV-2 spike-specific T cells, we performed surface and intracellular cytokine staining of CD4 and CD8 T cells as described previously.^11, 12^ Briefly, PBMCs were incubated in 1 mL RPMI 1640 medium containing 50 U/mL benzonase nuclease (Millipore, Darmstadt, Germany), 10% FBS, and penicillin–streptomycin for 1 h. Next, the cells were incubated in 200 µL medium with or without peptides (17-mers overlapping by 10 residues) corresponding to the full-length SARS-CoV-2 spike (Wuhan-1, BA.1, or BA.5), at a final concentration of 2 µg/mL of each peptide, for 30 min. Thereafter, 0.2 µL BD GolgiPlug and 0.14 µL BD GolgiStop (both from BD Biosciences) were added and incubated for 5.5 h. The cells were then stained using a LIVE/DEAD Fixable Blue Dead Cell Stain Kit (Thermo Fisher Scientific) and anti-CD3 (SP34-2), anti-CD8 (RPA-T8), anti-CD4 (L200), anti-CD45RO (UCHL1), and anti-CD27 (O323) antibodies. After fixation and permeabilization using the Cytofix/Cytoperm kit (BD Biosciences), the cells were stained with anti-CD154 (TRAP1) and anti-IFN-g (4S.B3), anti-TNF (MAb11), anti-IL-2 (MQ-17H12), anti-IL-4 (8D4-8), and anti-IL-13 (JES10-5A2). The cells were analyzed using a BD FACSymphony A5 flow cytometer (BD Biosciences). Data were analyzed using FlowJo v. 10.8.1. After gating live single T cells based on the forward scatter area and height (FSC-A and -H), side scatter area (SSC-A), live/dead cell exclusion, and CD3 staining, we separated live single T cells into CD4+ and CD8+ T cells. Subsequently, CD4+ and CD8+ T cells were further divided into memory phenotypes based on the expression of their CD27 and CD45RO expression levels. We defined CD154+CD4+ T cells expressing IFN-g, TNF, or IL-2 as Th1 cells and those expressing IL4 or IL-13 as Th2 cells.^12–14^ For spike-specific CD8+ T cells, memory cells were gated based on IFN-g or TNF expression. The frequency of cytokine production was determined by background subtracting the DMSO-supplemented group as a control.

### 2.8 Statistical Analysis

Categorical variables, presented as numbers and percentages, were compared using the Chi-square test, and continuous variables, presented as medians with ranges, were compared using the Mann-Whitney U test. Anti-SARS-CoV-2 spike protein antibody titers, neutralizing antibody titers, and the percentage of SARS-CoV-2 spike-specific T cells in both groups were compared using the Mann-Whitney U test. Anti-SARS-CoV-2 spike protein and neutralizing antibody titers after the third and bivalent omicron-containing booster vaccine were compared using the Wilcoxon signed-rank test. The Mann-Whitney U test was used to analyze the changes in anti-SARS-CoV-2 spike protein antibody titers and neutralizing antibody titers in both groups after the third and bivalent vaccines. Statistical significance was set at P < 0.05. GraphPad Prism 8.0.0 (GraphPad Software Inc., San Diego, CA, USA) was used for all analyses.

## 3. Result

### 3.1 Patient characteristics

The study included 73 KTX recipients and 17 healthy controls who received up to third doses of the SARS-CoV-2 mRNA vaccine. Characteristics of the KTX recipients and healthy controls are presented in Table 1. The median ages of KTX recipients and healthy controls were 59 (range: 28–73) and 35 (range: 23–67) years, respectively (p = 0.0025). The median durations after the third vaccination in the KTX recipients and healthy controls were 5.7 (range: 2.1–7.9) and 7.4 (range: 2.3–11.3) months, respectively (p < 0.0001).

**Table 1.**
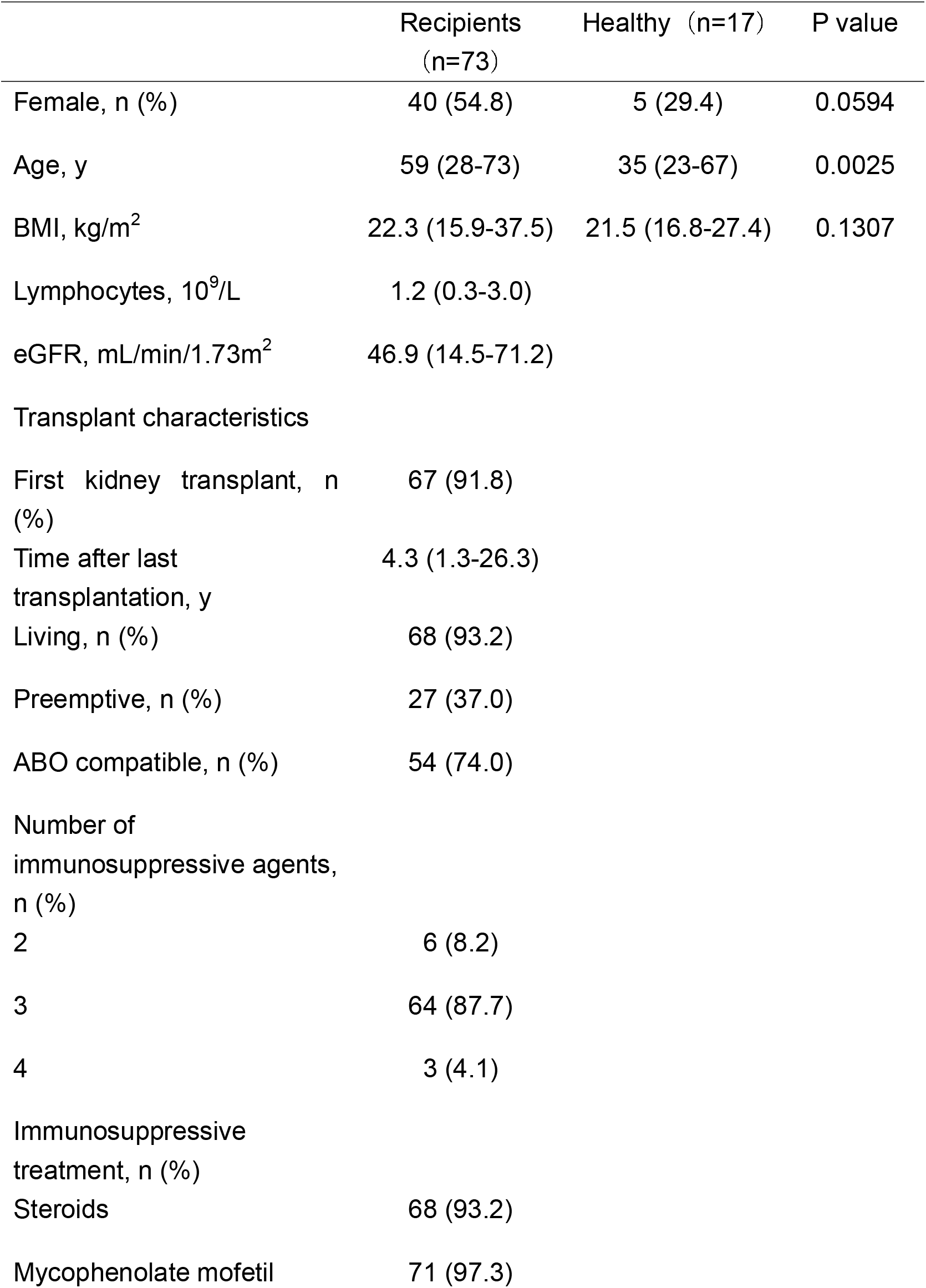
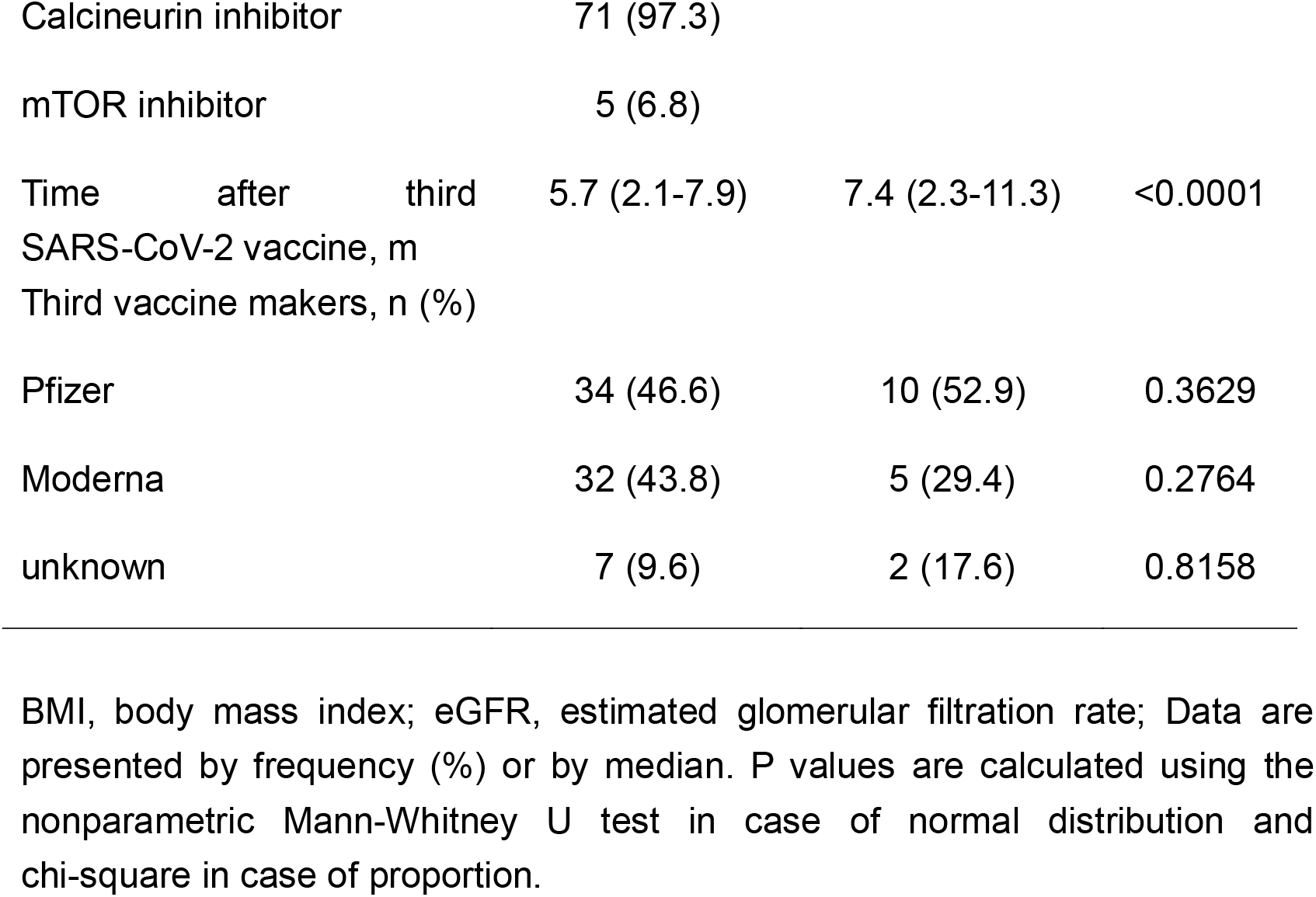
Baseline characteristics of kidney transplant recipients and healthy controls.

### 3.2 Anti-SARS-CoV-2 spike protein IgG antibody production by SARS-CoV-2 vaccination in KTX recipients

Anti-SARS-CoV-2 spike protein IgG antibody titers in both groups are shown in Figure 1. The median antibody titers in the KTX recipients and healthy controls after the third doses of vaccine were 7.8 (range: <1–2540) and 143.0 (range: 38.5–1370) AU/mL respectively, and the antibody titers in the KTX recipients were significantly lower than those in healthy controls (p < 0.0001). The antibody positivity rates were 67.1% and 100%. The background factors for the responders and non-responders among KTX recipients are summarized in Table 2. The time after KTX was significantly longer in the responder group than in the non-responder group (p=0.0012). Furthermore, The Mycophenolic acid-area under the plasm concentration-time curve levels were significantly lower in responders than in non-responders (p=0.0041). No significant differences were observed in the other background factors.

**Figure 1.**
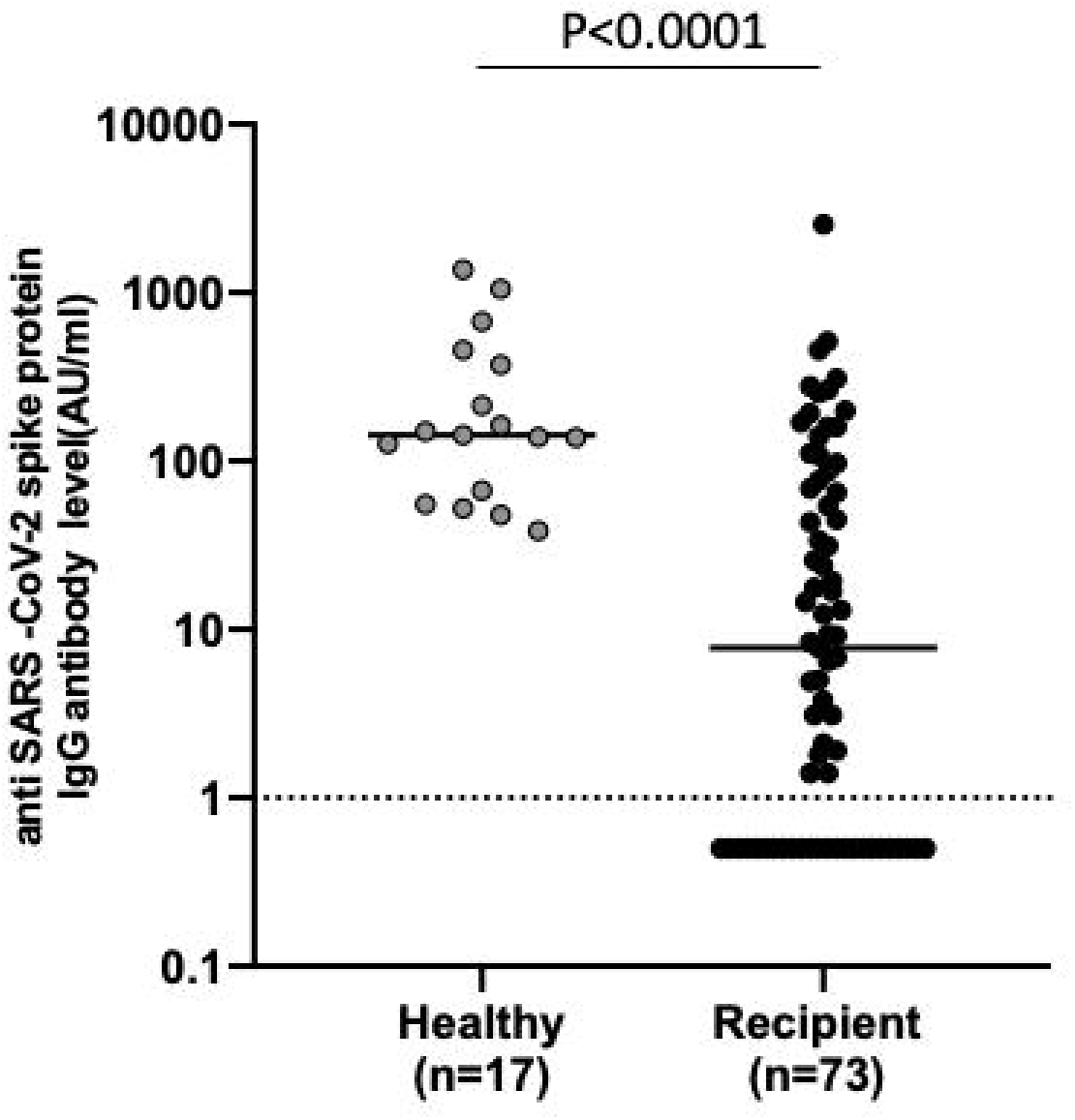
Anti-SARS-CoV-2 spike protein IgG antibody titers in healthy controls and KTX recipients. Anti-SARS-CoV-2 spike protein IgG antibody titers in 17 healthy controls and 73 KTX recipients. The cut-off value was defined as 1.0 AU/mL. P-values were calculated using the nonparametric Mann-Whitney U test.

**Table 2.**
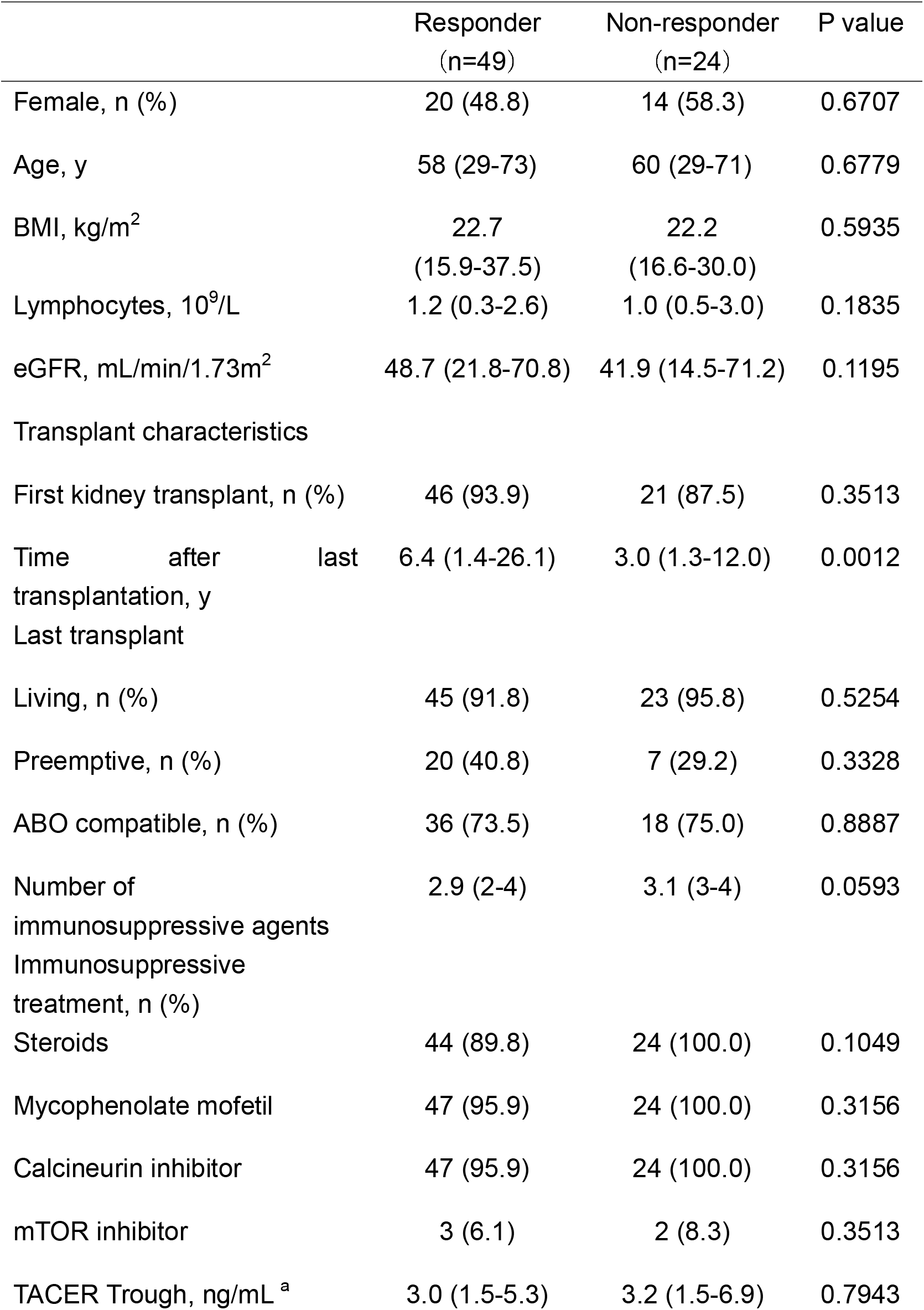
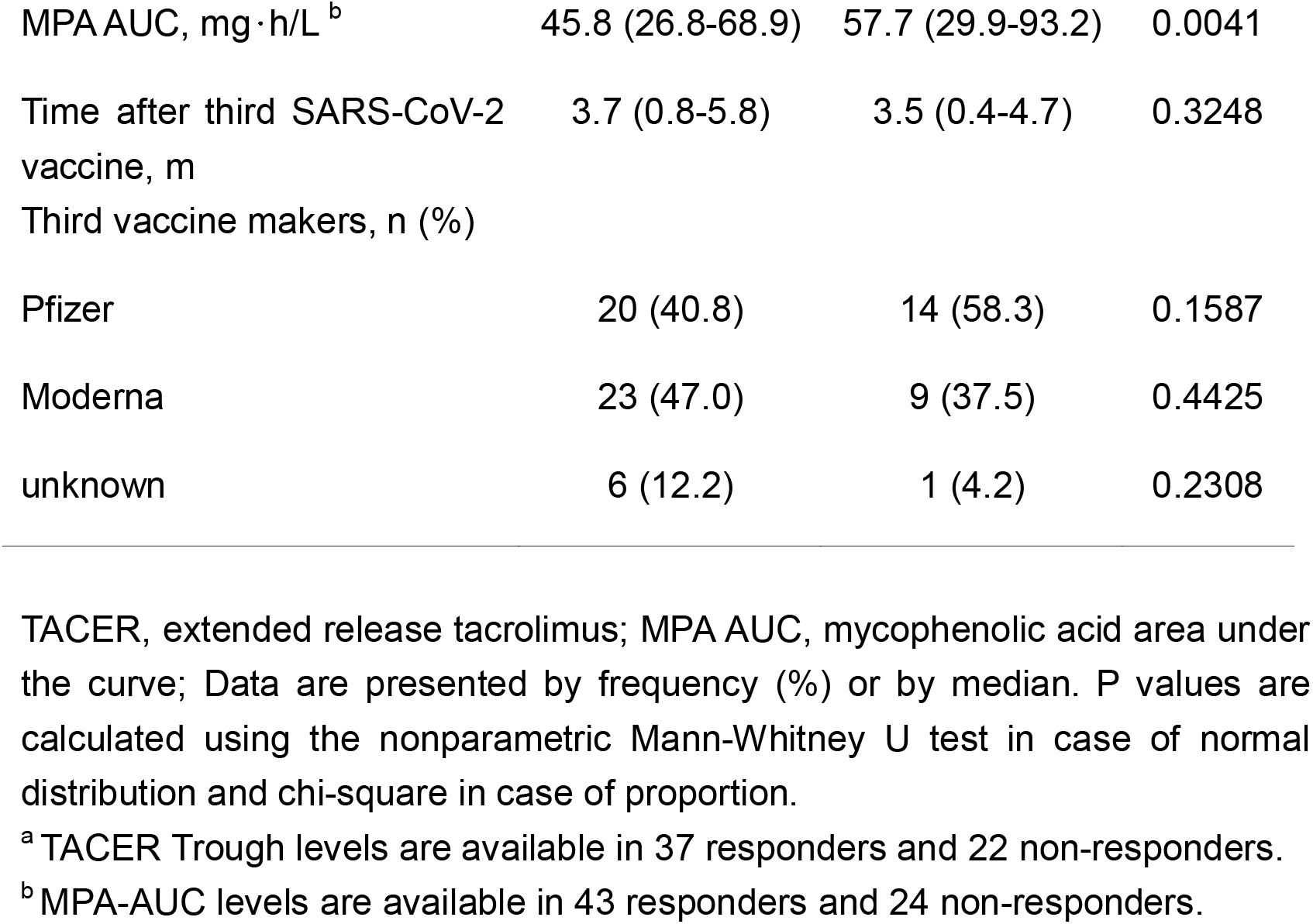
Differences in subject characteristics between Responder vs Non-responder in kidney transplant recipients.

### 3.3 Neutralizing activity of sera against SARS-CoV-2 variants after the third vaccine dose

To measure the neutralizing activity of sera from KTX recipients against B.1.1(WT), BA.1, and BA.5, we generated GFP-carrying recombinant SARS-CoV-2 with spike protein of WT, BA.1, or BA.5 by reverse genetics (rWT S-GFP, rBA.1 S-GFP, and rBA.5 S-GFP, respectively) (Figure 2a). Using these chimeric recombinant viruses, we performed a high-throughput neutralization assay evaluated by visual examination of the GFP signal. We previously compared the 50% neutralizing titer (NT_50_) calculated by reverse transcription quantitative polymerase chain reaction (RT-qPCR) and neutralizing activity titer calculated by visual measurement of GFP expression using a fluorescent microscope. The NT_50_ calculated from viral RNA levels in the supernatants of infected cells were found to be well correlated with the titer of neutralizing antibody calculated by GFP fluorescence intensity.^15^ The neutralizing antibody titers against rWT S-GFP, rBA.1 S-GFP, and rBA.5 S-GFP in KTX recipients and healthy controls after the third dose of the mRNA vaccine are shown in Figure 2b. The neutralizing antibody titers against all types of viruses in KTX recipients were significantly lower than those in healthy controls (p < 0.0001). Interestingly, when the virus was mutated, the neutralizing antibody titers in both groups were significantly decreased (Figure 2c and d, p < 0.0001). These results suggest that rBA.1 and rBA.5 S-GFP evade neutralizing antibodies not only in healthy controls but also in KTX recipients.

**Figure 2.**
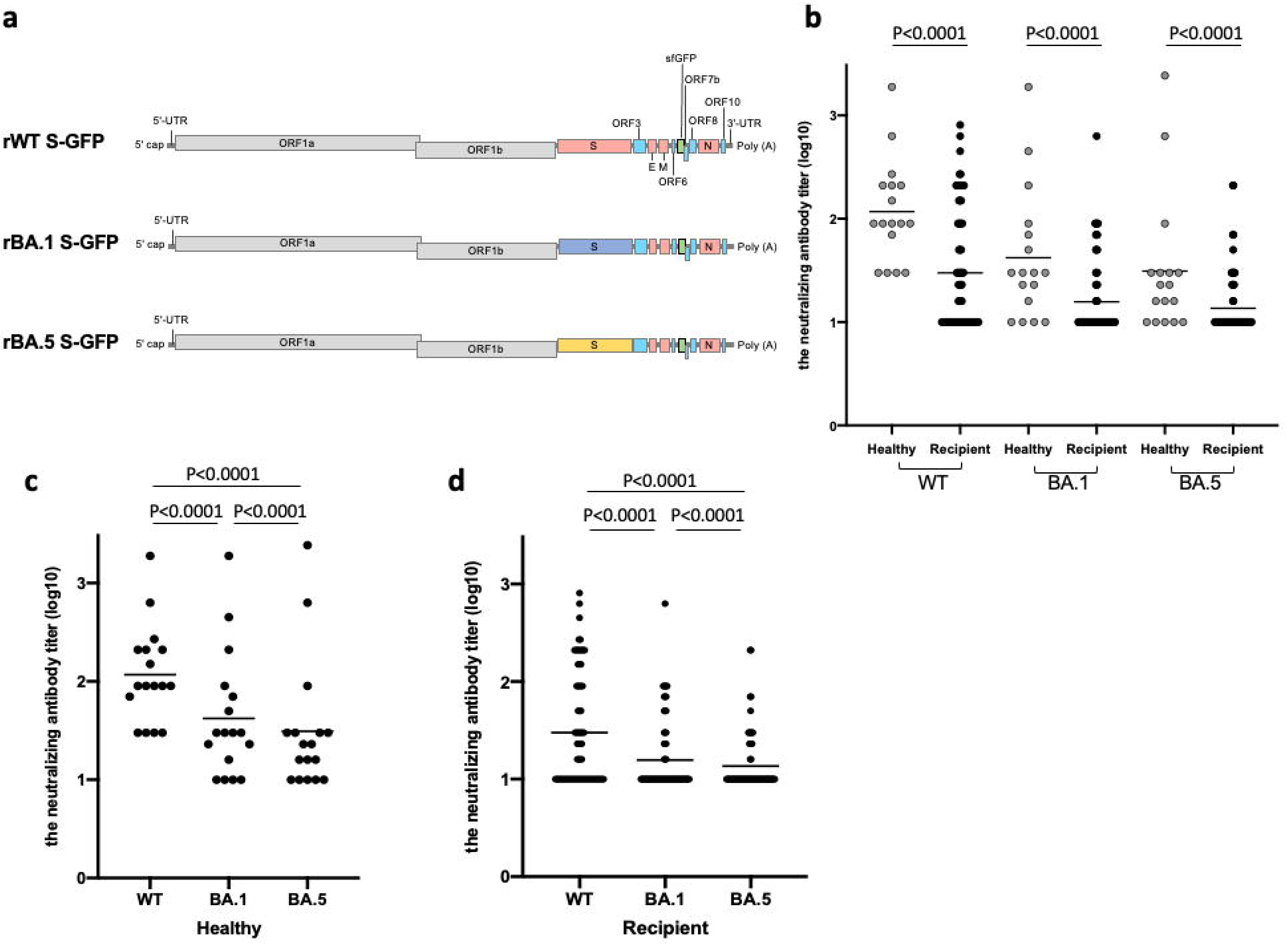
Neutralizing activity against SARS-CoV-2 variants in healthy controls and KTX recipients after third vaccine dose. (A) We generated GFP-carrying recombinant SARS-CoV-2 with spike protein of ancestral B1.1 (WT), BA.1, or BA.5 by reverse genetics (rWT S-GFP, rBA.1 S-GFP, and rBA.5 S-GFP, respectively). (B) The neutralizing antibody titer (log10) against rWT S-GFP, rBA.1 S-GFP, and rBA.5 S-GFP in healthy controls and KTX recipients after the third dose of mRNA vaccine. The mixture of chimeric recombinant SARS-CoV-2 and diluted serum was inoculated into VeroE6/TMPRSS2 cells. After 36 hours post-infection, the expression of GFP in VeroE6/TMPRSS2 cells was observed by florescent microscopy and the titer of neutralizing antibody was calculated. P-values were calculated using the nonparametric Mann-Whitney U test. (C) The neutralizing antibody titer (log10) against rWT S-GFP, rBA.1 S-GFP, and rBA.5 S-GFP in healthy controls. This data was extracted from (B). P-values were calculated using the Wilcoxon signed-rank test. (D) The neutralizing antibody titer (log10) against rWT S-GFP, rBA.1 S-GFP, and rBA.5 S-GFP in KTX recipients. This data was extracted from (B). P-values were calculated using the Wilcoxon signed-rank test.

### 3.4 Induction of spike-specific T-cell responses in healthy donors and KTX recipients

In general, immunosuppressive drugs, including calcineurin inhibitors, suppress T cell activity. To investigate whether spike-specific CD4+ T cell responses in KTX recipients were induced by mRNA vaccines, we performed flow cytometry analysis. PBMCs obtained from healthy donors and KTX recipients were stimulated with overlapping peptides corresponding to the SARS-CoV-2 spike protein. We defined CD154+CD4+ T cells expressing IFN-g, TNF, or IL-2 as Th1 cells and those expressing IL4 or IL-13 as Th2 cells (Figure 3a).^12–14^ Spike-specific Th1-cell and Th2-cell frequencies were determined in the stimulated samples by subtracting the background signal observed in the unstimulated samples. The frequency of spike-specific Th1 cells against WT spikes in KTX recipients was comparable to that in healthy controls (Figure 3b, p = 0.7314). Furthermore, mRNA vaccine induced spike-specific Th1 CD4 T-cell responses against BA.1 and BA.5 were at comparable levels to those of Th1 CD4 T cells responses against WT in both healthy controls and KTX recipients. In contrast, the frequency of spike-specific Th2 cells against WT, BA.1, and BA.5 spikes in KTX recipients was higher than that in healthy controls (Figure 3c, WT: p = 0.0312, BA.1: p = 0.0037, BA.5: p = 0.0474). These results suggest that mRNA vaccines can induce spike-specific CD4+ T-cell responses to omicron sublineages not only in healthy controls but also in KTX recipients. Besides, KTX recipients are more susceptible to the induction of Th2-biased CD4+ T cell responses.

**Figure 3.**
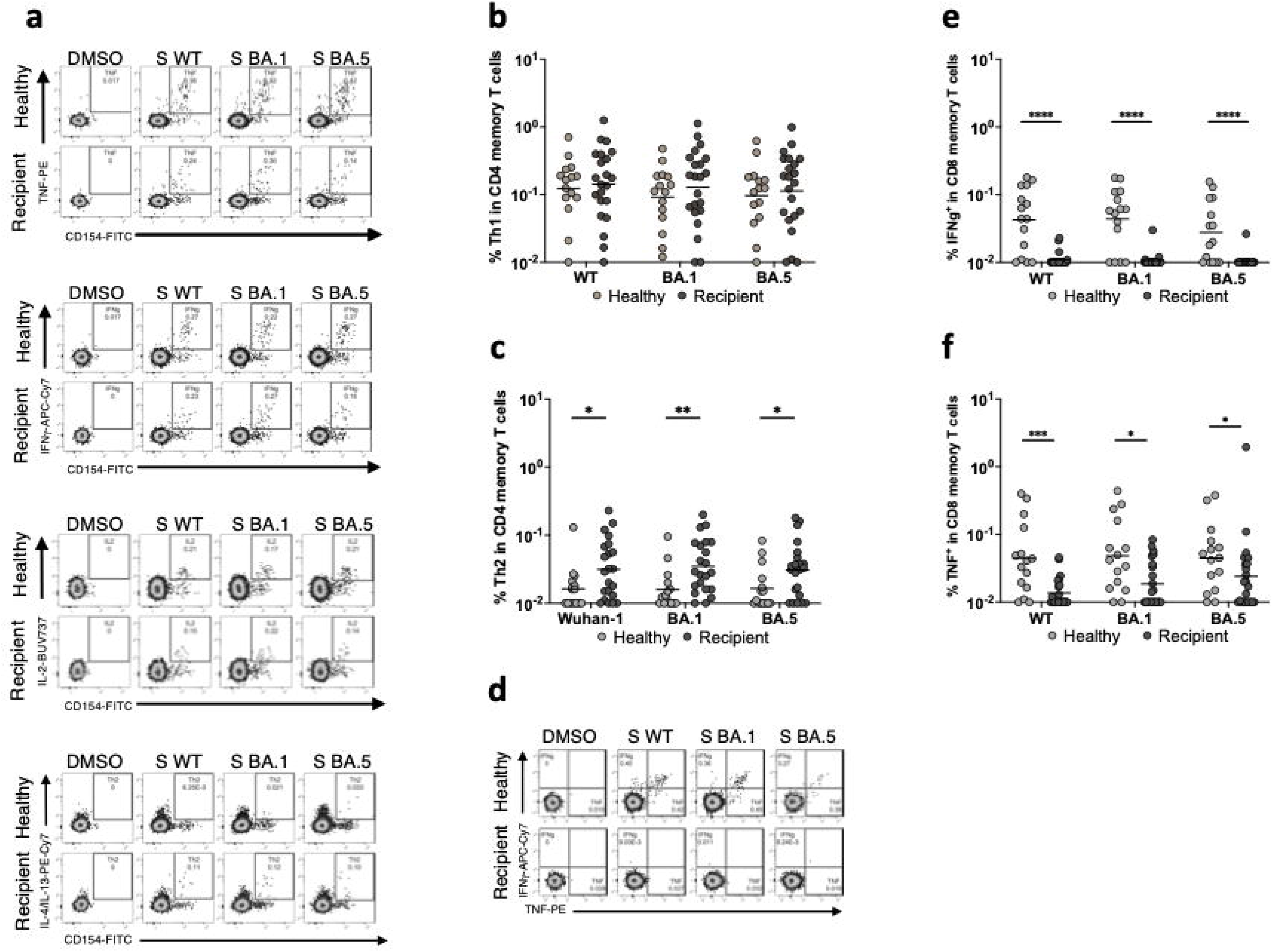
Spike-specific T-cell responses in healthy controls and KTX recipients. (A) After gating live single T cells, based on forward scatter area and height (FSC-A and -H), side scatter area (SSC-A), live/dead cell exclusion, and CD3 staining, we separated the PBMCs into CD4+ and CD8+ T cells. Subsequently, CD4+ and CD8+ T cells were further divided into memory phenotypes based on the expression of CD27 and CD45RO. We defined CD154+CD4 T cells expressing IFN-γ (upper left panels), TNF (upper right panels), or IL-2 (bottom left panels) as Th1 cells and expressing IL4 or IL-13 (bottom right panels) as Th2 cells. (B) The frequency of spike-specific Th1 CD4 T cells against Wuhan-1 (Healthy controls; 0.01%-0.704%, geometric mean = 0.123%, KTX recipients; 0.01%-1.254%, geometric mean = 0.142%), BA.1 (Healthy controls; 0.012%-0.517%, geometric mean = 0.097%, KTX recipients; 0.01%-1.123%, geometric mean = 0.128%), or BA.5 (Healthy controls; 0.01%-0.654%, geometric mean = 0.1%, KTX recipients; 0.01%-0.986%, geometric mean = 0.113%) in CD4 total memory T cells from vaccinated healthy controls and KTX recipients. P-values were calculated using the nonparametric Mann-Whitney U test. (C) The frequency of spike-specific Th2 CD4 T cells against Wuhan-1 (Healthy controls; 0.01%-0.13%, geometric mean = 0.016%, KTX recipients; 0.01%-0.23%, geometric mean = 0.032%), BA.1 (Healthy controls; 0.01%-0.095%, geometric mean = 0.016%, KTX recipients; 0.01%-0.2%, geometric mean = 0.035%), or BA.5 (Healthy controls; 0.01%-0.083%, geometric mean = 0.016%, KTX recipients; 0.01%-0.18%, geometric mean = 0.031%) in CD4 total memory T cells from vaccinated healthy controls and KTX recipients. P-values were calculated using the nonparametric Mann-Whitney U test. (D) After gating live single T cells, based on forward scatter area and height (FSC-A and -H), side scatter area (SSC-A), live/dead cell exclusion, and CD3 staining, we separated the PBMCs into CD4+ and CD8+ T cells. Subsequently, CD4+ and CD8+ T cells were further divided into memory phenotypes based on the expression of CD27 and CD45RO. For spike-specific CD8 T cells, memory cells were gated based on the expression of IFN-γ or TNF. (E) The frequency of spike-specific CD8 T cells producing IFN-γ against Wuhan-1 (Healthy controls; 0.01%-0.18%, geometric mean = 0.042%, KTX recipients; 0.01%-0.023%, geometric mean = 0.011%), BA.1 (Healthy controls; 0.01%-0.177%, geometric mean = 0.044%, KTX recipients; 0.01%-0.03%, geometric mean = 0.011%), or BA.5 (Healthy controls; 0.01%-0.155%, geometric mean = 0.028%, KTX recipients; 0.01%-0.026%, geometric mean = 0.01%) in CD8 total memory T cells from vaccinated healthy controls and KTX recipients. P-values were calculated using the nonparametric Mann-Whitney U test. (F) The frequency of spike-specific CD8 T cells producing TNF against Wuhan-1 (Healthy controls; 0.01%-0.4%, geometric mean = 0.044%, KTX recipients; 0.01%-0.046%, geometric mean = 0.014%), BA.1 (Healthy controls; 0.01%-0.44%, geometric mean = 0.048%, KTX recipients; 0.01%-0.084%, geometric mean = 0.019%), or BA.5 (Healthy controls; 0.01%-0.38%, geometric mean = 0.045%, KTX recipients; 0.01%-1.95%, geometric mean = 0.024%) in CD8 total memory T cells from vaccinated healthy controls and KTX recipients. P-values were calculated using the nonparametric Mann-Whitney U test.

In addition to antibodies and CD4+ T-cell responses, spike-specific CD8+ T-cell responses also contribute to defense against SARS-CoV-2 infection^11, 16–18^. We next examined whether CD8+ T cell responses were induced by mRNA vaccines in KTX recipients using flow cytometry. Spike-specific CD8+ T cells were gated based on the expression of IFN-g or TNF in total memory CD8+ T cells (Figure 3d). The frequencies of spike-specific CD8+ T cells producing IFN-g and TNF against WT, BA.1, and BA.5 spikes in KTX recipients were lower than those in healthy controls (Figure 3e, WT: p < 0.0001, BA.1: p < 0.0001, BA.5: p < 0.0001 and Figure 4f, WT: p = 0.0005, BA.1: p = 0.011, BA.5: p = 0.0405). These results suggest that immunosuppressive drugs potentially affect spike-specific CD8+ T-cell responses in KTX recipients.

**Figure 4.**
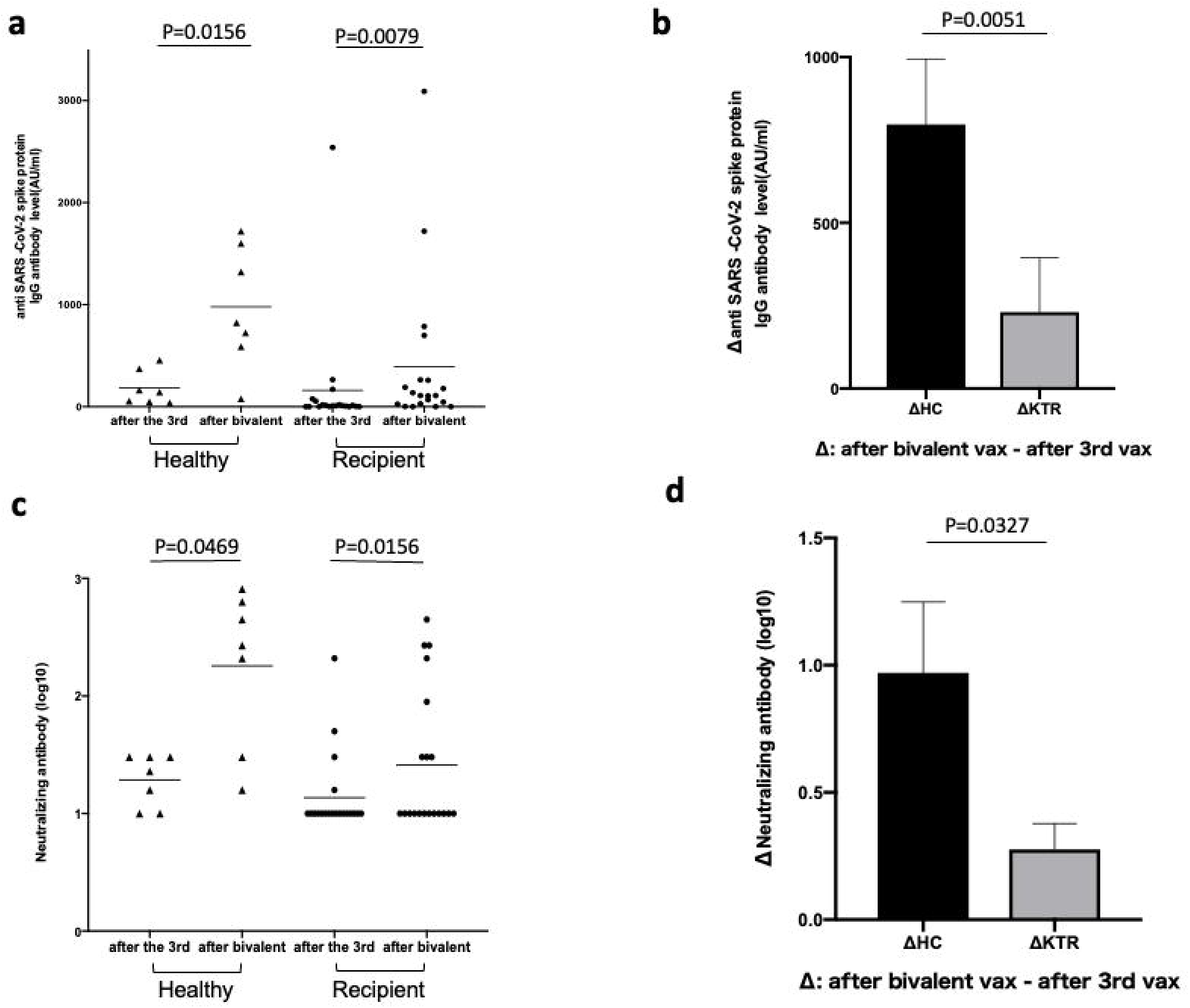
Anti-SARS-CoV-2 spike protein IgG antibody titers and neutralizing activity against rBA.5 S-GFP virus in healthy controls and KTX recipients after a third vaccine dose and bivalent omicron-containing booster vaccine. (A) Anti-SARS-CoV-2 spike protein IgG antibody titers in healthy controls and KTX recipients after a third vaccine dose and bivalent omicron-containing booster vaccine. P-values were calculated using the Wilcoxon signed-rank test. (B) The increase in anti-SARS-CoV-2 spike protein IgG antibody titers in healthy controls and KTX recipients after bivalent omicron-containing booster vaccine. P-values were calculated using the nonparametric Mann-Whitney U test. (C) The neutralizing antibody titer (log10) against rBA.5 S-GFP virus in healthy controls and KTX recipients after a third vaccine dose and bivalent omicron-containing booster vaccine. The mixture of rBA.5 S-GFP virus and diluted serum of healthy controls or KTX recipients was inoculated into VeroE6/TMPRSS2 cells. After 36 hours post-infection, the expression of GFP in VeroE6/TMPRSS2 cells was observed by florescent microscopy and the titer of neutralizing antibody was calculated. P-values were calculated using the Wilcoxon signed-rank test. (D) The increase in the neutralizing antibody titer (log10) against rBA.5 S-GFP virus in healthy controls and KTX recipients after bivalent omicron-containing booster vaccine. P-values were calculated using the nonparametric Mann-Whitney U test.

### 3.5 Neutralizing activity of sera against rBA.5 S-GFP virus after the bivalent omicron-containing booster vaccine

Serum samples were collected from 20 KTX recipients and 7 healthy controls after administering the bivalent omicron-containing booster vaccine. Of the 20 KTX recipients who received the bivalent omicron-containing booster vaccine, three received the bivalent vaccine after the third dose and 17 received the bivalent vaccine after the fourth dose. All seven healthy controls received the bivalent omicron-containing booster vaccine after the third dose. We measured anti-SARS-CoV-2 spike protein IgG antibody titers and performed a neutralizing antibody assay using the rBA.5 S-GFP virus. The anti-SARS-CoV-2 spike protein IgG antibody titers and neutralizing activity of both groups are shown in Figure 5. Anti-SARS-CoV-2 spike protein IgG antibody titers were significantly elevated in both groups (Figure 4a, recipients: p = 0.0079, healthy controls: p = 0.0156). However, the increase in anti-SARS-CoV-2 spike protein IgG antibody titers in KTX recipients was lower than that in healthy controls (Figure 4b, p = 0.0051). Similarly, the neutralizing activity in both groups was significantly elevated (Figure 4c, recipients: p = 0.0156, healthy controls: p = 0.0469); however, the increase in the neutralizing antibody titer in KTX recipients was smaller (Figure 4d, p = 0.0327). These results suggest that the Anti-SARS-CoV-2 spike protein IgG antibody titers and neutralizing activity of KTX recipient’s sera vaccinated with the bivalent omicron-containing booster vaccine were not much higher than those of healthy controls.

## 4. Discussion

To the best of our knowledge, this is the first study to investigate the neutralizing antibody titers and T cell responses against SARS-CoV-2 Omicron BA.5 in the sera of KTX recipients after a third dose of an mRNA SARS-CoV-2 vaccine and a bivalent omicron-containing booster vaccine. We found that kidney recipients did not gain sufficient immunity against omicron BA.5 with the third dose of vaccine, and the effect of bivalent omicron-containing booster vaccine was also limited.

First, we evaluated the anti-SARS-CoV-2 spike protein IgG antibody and detected seropositivity in 67.1% of the KTX recipients after the third vaccination. Similarly, Louise et al. reported an antibody-positive rate of 71% in KTX recipients three weeks after the third vaccination.^6^ Al Jurdi et al. reported that the percentage of KTX recipients with anti-SARS-CoV-2 spike protein IgG antibodies increased from 29% to 67% after a third vaccination.^19^ Thus, antibody titers were increased after the third vaccination but were considerably lower than those in healthy controls. We showed that one predictor of non-response among KTX recipients was a shorter time after kidney transplantation. The higher risk may be primarily attributed to higher levels of immunosuppression early on, shortly after KTX. We also found that a high mycophenolate mofetil (MMF) blood concentration was significantly associated with the risk of being a non-responder. In contrast, the tacrolimus trough concentration was similar between the two groups. Miura et al. reported that IgG positivity is significantly associated with MMF cessation or dose reduction.^20^ Tomiyama et al. reported that seropositivity was 95% after the third dose of the vaccine in liver transplant recipients, only 48% of whom were taking MMF.^15^ These results suggest that MMF strongly contributes to the suppression of antibody production in KTX recipients.

We showed that the neutralizing titers against Omicron BA.5 in KTX recipients were significantly lower than those in healthy controls using the live SARS-CoV-2 virus neutralization assay, suggesting that recipients are at a higher risk of infection and that the vaccine is less effective in KTX recipients. Tuekprakhon et al. reported that as the mutation progressed, neutralizing titers decreased in the general population after a third dose of the vaccine.^21^ We also showed that the neutralizing titers also decreased in KTX recipients as the virus showed mutations from WT to BA.1 to BA.5, suggesting that recipients are at a higher risk of infection as the mutation progresses.

T-cell responses have been shown to contribute to the reduction of severe illnesses caused by SARS-CoV-2. KTX recipients are at a risk of severe SARS-CoV-2 infection because of their reduced T cell immunity.^22^ A previous study showed that the third mRNA dose induced a T cell response against WT spike peptides in KTX recipients, and cellular immunity to Omicron BA.1 variant immunogenic peptides was low in both KTX recipients and healthy controls.^8^ However, no studies have evaluated the T cell response to Omicron BA.5 spike peptides. We found the frequency of spike-specific Th1 cells against BA.5 spike in KTX recipients was comparable to that in healthy controls; however, Th2 CD4+ T cell responses were significantly higher in KTX recipients. The Th1/Th2 balance of CD4+ T cells is important in cellular responses, and Th2 cells-skewed CD4+ T cells cause an increased risk of severe infection and vaccine-associated enhanced respiratory disease.^23, 24^ Further investigation will be needed to evaluate the potential risks of increasing TH2 CD4+ T-cell responses in KTX.

We also showed that the number of spike-specific CD8+ T cells producing IFN-g and TNF was significantly lower in KTX recipients than in healthy controls. These results suggest that KTX recipients did not acquire sufficient immunity against Omicron BA.5 (neither humoral immunity nor in cellular immunity), even after receiving three doses of the vaccine.^17, 23, 25^ Furthermore, we evaluated the effects of a bivalent Omicron-containing booster vaccine in KTX recipients. Although the current bivalent Omicron-containing booster vaccine produced increased anti-SARS-CoV-2 spike protein IgG and neutralizing antibody titers, these effects were lower in KTX recipients than in healthy controls. Therefore, although the severity the Omicron variants is less compared to the previous lineages,^26, 27^ KTX recipients might be more susceptible to Omicron BA.5 and develop more severe disease.

French groups reported the efficacy of tixagevimab/cilgavimab in reducing the morbidity and severity of SARS-CoV-2 in kidney recipients who failed to develop a protective humoral response after at least three doses of an mRNA vaccine during the omicron BA.1-BA.2 epidemic.^28, 29^ On the other hand, in vitro data showed a substantial reduction in the activity of tixagevimab/cilgavimab against omicron BA.4/5, BQ.1.1, and XBB.1.5. ^30^ Although the efficacy of tixagevimab/cilgavimab is different among omicron variants, these reports suggested that tixagevimab/cilgavimab administration is an alternative approach for kidney recipients. Recently, Solera et al. reported that tixagevimab/cilgavimab improved the neutralization of BA.4/5, but this effect was not observed against BQ.1.1 and XBB.1.5.^31^ Therefore, new effective neutralizing antibodies against BQ.1.1, XBB.1.5, and new variants to emerge in the future must be developed for KTX recipients.

The strength of our study is the analysis of neutralizing activity using the live SARS-CoV-2 virus neutralization assay. Furthermore, this is the first study investigating T cell responses against SARS-CoV-2 Omicron BA.5 and the effect of bivalent Omicron-containing booster vaccine in KTX recipients. Nonetheless, there are some limitations of our study. First, the time between vaccination and blood collection varied because the blood samples were collected during outpatient visits. Although KTX recipients had a shorter term than controls, KTX recipients had lower neutralizing activity. Therefore, the timing of sample collection did not appear to have a significant effect on the results. Second, our cohort lacked an age-matched control group and had a small sample size. In conclusion, the adaptive immunity of KTX recipients was not significantly developed after a third dose of an mRNA vaccine, suggesting that recipients are at a higher risk for infection and severe disease than healthy controls. The efficacy of bivalent omicron-containing booster vaccines is poorer in recipients than in healthy controls; therefor, other methods need to be explored.

## Abbreviations

CLEIA: chemiluminescent enzyme immunoassay
CPER: circular polymerase extension reaction
DMEM: Dulbecco’s modified Eagle’s medium
FBS: fetal bovine serum
GFP: green fluorescent protein
KTX: kidney transplant
MMF: mycophenolate mofetil
PBMC: peripheral blood mononuclear cell
SARS-CoV-2: severe acute respiratory syndrome coronavirus 2
TCID50: 50% tissue culture infective dose
WT: Wuhan-type

## Acknowledgments

We would like to thank Editage (https://www.editage.jp/) for English language editing.

## Data availability statement

All data generated or analyzed during this study are included in this article. Further enquiries can be directed to the corresponding author.

## Supporting information statement

Additional supporting information may be found online in the Supporting Information section.

